# Coronavirus Spike-RBD Variants Differentially Bind to the Human ACE2 Receptor

**DOI:** 10.1101/2024.03.07.583944

**Authors:** Paul Feinstein

**Affiliations:** Department of Biological Sciences, Hunter College, City University of New York, New York, NY 10065; The Graduate Center Programs in Biochemistry, Biology and CUNY Neuroscience Collaborative, 365 5th Ave, New York, NY 10016

**Author notes:** Correspondence: Paul Feinstein, 695 Park Avenue, 904 North Building, Department of Biology, New York, NY 10065. Contributions: P.F. conceived the project. P.F. designed the experiments. P.F. performed all experiments. P.F. analyzed the data. P.F. wrote the manuscript.

**Keywords:** SARS-CoV-2, Spike protein, RBD, hACE2, Omicron variant, Xbb.1.5 variant, NL63, ACE2 downregulation

## Abstract

The SARS-CoV-2 betacoronavirus infects people through binding the human Angiotensin Receptor 2 (ACE2), followed by import into a cell utilizing the Transmembrane Protease, Serine 2 (TMPRSS2) and Furin cofactors. Analysis of the SARS-CoV-2 extracellular spike protein has suggested critical amino acids necessary for binding within a 197-residue portion, the receptor binding domain (RBD). A cell-based assay between a membrane tethered RBD-GFP fusion protein and the membrane bound ACE2-Cherry fusion protein allowed for mutational intersection of both RBD and ACE2 proteins. Data shows Omicron BA.1 and BA.2 variants have altered dependency on the amino terminus of ACE2 protein and suggests multiple epitopes on both proteins stabilize their interactions at the Nt and internal region of ACE2. In contrast, the H-CoV-NL63 RBD is only dependent on the ACE2 internal region for binding. A peptide inhibitor approach to this internal region thus far have failed to block binding of RBDs to ACE2, suggesting that several binding regions on ACE2 are sufficient to allow functional interactions. In sum, the RBD binding surface of ACE2 appears relatively fluid and amenable to bind a range of novel variants.

## Introduction

Since the 2019 SARS2-CoV-2 outbreak that has infected the global population, we are constantly being infected by novel mutations in the Spike protein as well as other portions of the virus. Due to vaccine protocols and drugs such as Paxlovid (1), infections once they happen yield outcomes with significantly reduced morbidities. In neither scenario is infection prevented from being initiated (2). Masks, when worn properly, mitigate some of the SARS2 infectivity, but the vast majority of individuals will not embrace this as a long-term solution to preventing infections. In additions, SARS2 is not the only virus that infects humans and can produce death. As such it is critical that the medical industry find viral transmission mitigation strategies (3–6). Transmembrane Protease, Serine 2 (TMPRSS2) is necessary to promote cleavage of the Spike S1 protein affording the virus enhanced transmission through binding to the multimeric human Angiotensin-Converting Enzyme 2 protein, ACE2. Small molecules, although not volatile, that may be able to limit viral load has been identified, which targets TMPRSS2 functionality (3).

The two betacoronavirus’ SARS-CoV-1 (SARS1), SARS-CoV-2 (SARS2) as well as the alphacoronavirus H-CoV-NL63 (NL63) all bind ACE2, expressed by Sustentacular cells in the nasal olfactory epithelium, to initiate human infection (7–9). The Spike protein on coronavirus’ coat establishes this ACE2 binding through a short stretch of residues referred to as the Receptor Binding Domain (RBD) portion: 196 residues (SARS1) (10) ; 197 residues (SARS2) and (135 residues) NL63. Given that two evolutionarily distinct coronaviruses, RBDs SARS2 and NL63 share 7% amino acid identity, attack the same target protein (ACE2), it seems likely that the biological properties of ACE2 make it an attractive target for cellular entry. We have previously described an *in vitro* approach to study the interactions between an amino terminal membrane-tethered RBD fused to the β2 adrenergic receptor (β2AR)-GFP and an ACE2 protein fused to mCherry (See Feinstein 2024a). In this assay, the RBDs for SARS1, SARS2 and NL63 cause rapid downregulation and internalization of ACE2, even without expression of other viral fusion polypeptides that are unmasked during coronavirus infection (See Feinstein 2024a).

We have shown disulfide bond denaturing agents can abolish RBD functionality *in vitro* and still require additional *in vivo* testing. To further pursue how RBD binding to ACE2 occurs we undertook a structure-function mutagenesis on ACE2, to identify an antagonist of binding. The ACE2 protein initially traffics to the endoplasmic reticulum via a signal peptide that is eventually cleaved, producing a single pass plasma membrane mature ACE2 protein, residues 19-805. Crystallographic studies describing SARS1-RBD and SARS2-RBD binding to ACE2 revealed sixteen residues distributed within the amino terminal (Nt) 19-83 residues, as well as another five residues within an internal region containing residues 330, 353-357.

SARS2-RBD residues necessary for ACE2 binding were previously characterized in a viral-free assay, revealing the distributed nature of interactions (Feinstein 2024a). One key amino acid, Y489A juxtaposed to the C480-C488 disulfide linkage (disulfide Loop 4) of the SARS2-RBD that did not affect plasma membrane trafficking of the protein, but nearly abolished the interactions, which pointed to an important domain in the ACE2 interacting region within the receptor binding motif (RBM). Mutations around disulfide Loop 4 within SARS2-RBD that reduced or abolished binding to ACE2, suggested ACE2 Nt proximal regions that might be more critical for binding.

Here we show mutagenesis of the ACE2 Nt region identifies a swap of eight residues of *R. Pusillus* into the Nt proximal portion of the human ACE2 (residues between 20-34), which was able to abolish SARS2-RBD binding. But both Omicron BA.1 and BA.2-RBDs were still able to interact with this ACE2 mutant. The addition of three mutations (L79A, M82A, Y83A-LMY➔A) in the Nt distal region of ACE2 were able to completely abolish Omicron BA.1 or BA.2-RBD internalization, while the control NL63-RBD from alphacoronavirus was not affected by any Nt proximal or distal mutations. Both NL63-RBD and SARS2-RBD were both unable to bind ACE2 when its internal residues K353 and D355 are mutated. Using co-crystallization data as a guide, a double mutant T500A, G502A on SARS2-RBD is also able to abolish its binding. A peptide based on K353-G354-D355 of ACE2 was not able to inhibit either SARS2-RBD nor NL63-RBD binding to occur. The totality of the data suggest: 1) multiple weak binding sites on ACE2 sum to create high affinity; 2) the internal region residues may be more universal RBD binding sites necessary for internalization; and 3) there are several permutations of RBD binding to ACE2. The changing nature of RBD variants binding preferentially to different ACE2 residues may correlate with the severity of symptoms by altering the degree of internalization by the ACE2 protein.

## Results

### ACE2 Amino terminal domain mutagenesis

SARS2-RBD is predicted to bind a number of residues on ACE2. These interaction domains can be separated into Nt proximal region: 19-45; Nt distal region: 79, 82-83, Internal region 1: 330, and Internal region 2: 353-357. Both RaTG13 and SARS1 RBDs are predicted to bind the same three regions with some variation in specific residues **(Figure 1A)**. The ACE2 Nt proximal region is poorly conserved (<50% homology) across 408 species from residues 19-27 but becomes increasingly more conserved from residues 28-45 (>50% homology)(11). By contrast, the ACE2 Nt distal region retains conservation only in residue Y83 (>50%) and the Internal region residues of 330 and 353-357 are also highly conserved (>50%) (**Figure 1A, B**). NL63-RBD has reduced contacts with the divergent Nt proximal region, no contacts with the divergent Nt distal region, and expanded contacts with the conserved Internal region 1. The regional overlap in binding to ACE2 for NL63-RBD and SARS2-RBD (SARS1 and RaTG13 as well) is remarkable despite the near absence of protein homology (Feinstein 2024a). The disulfide Loop 4 region of SARS2 is strongly implicated to bind only a the ACE2 Nt proximal and distal regions (**Figure 1B)**. Mapping studies show that both N487 and Y489 are in involved in coordinating binding at both the Nt proximal and distal regions (**Figure 1B**), suggesting that disruption of this contacts might abolish binding (12).

**Figure 1:**
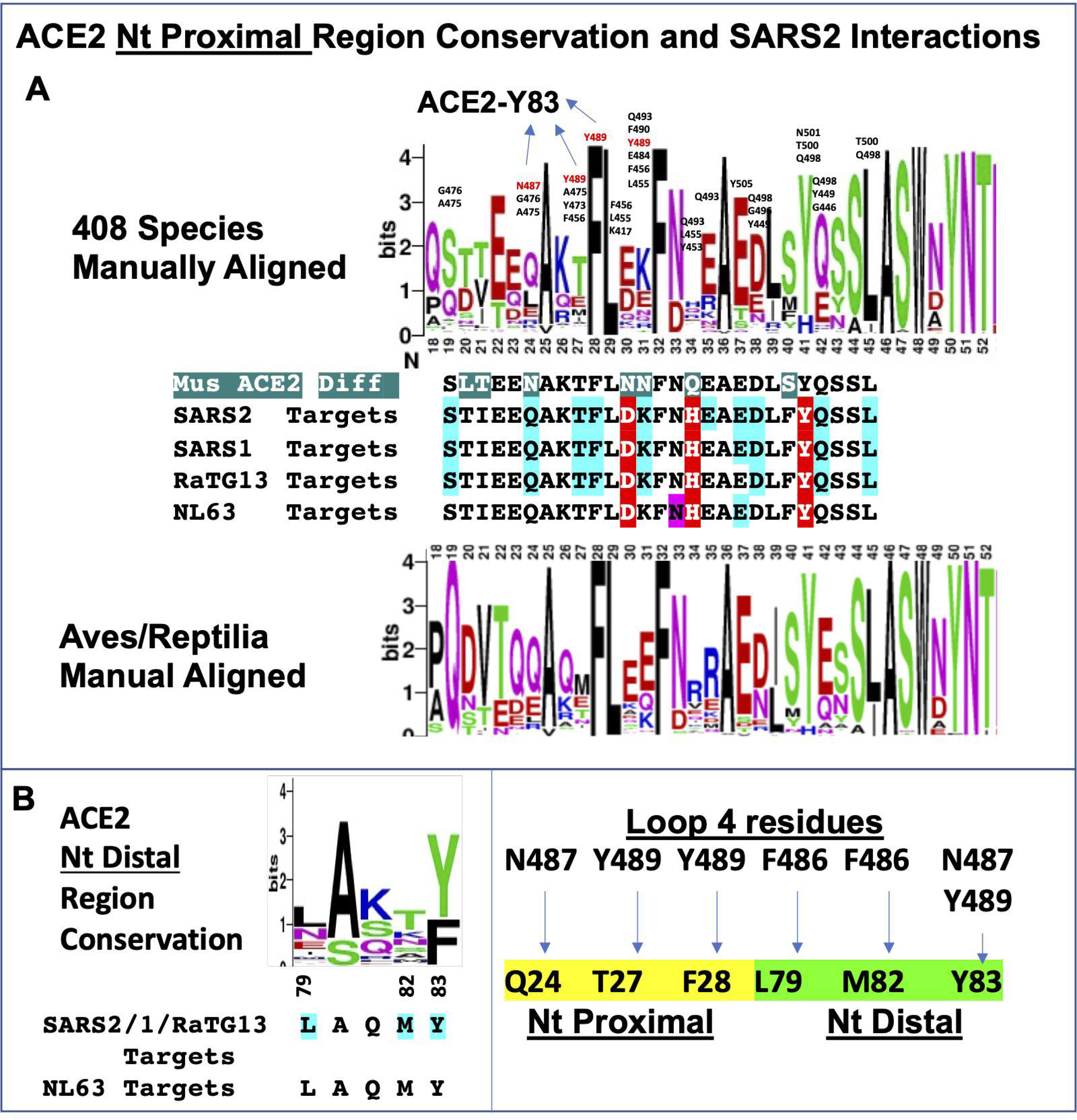
Conservation of ACE2 N-terminus and Predicted SARS2 Contact Sites. **A. ACE2 Nt. Proximal Region**. Logo plot of 408 ACE2 species (11) manually aligned starting from residue 18, the last residue before the signal peptide is spliced. ACE2 conserved residues between 22-41 appear to have a periodicity consistent with forming an alpha-helix whereby they would nominally be on the same face. SARS2 point of contacts are predominantly to the non-conserved ACE2 residues 19, 24, 27, 30, 31, 34, 35, 37, 38 and 42; conserved ACE2 residues F28, Y41 and L45. SARS2 contacting residues annotated within Logo plot. SARS2 N487 and Y489 residues in red also interact with ACE2-residue 83 (**See B**.). Logo plot of Aves/Reptilla is shown, which has a divergent amino terminus. Between Logo Plots reveals comparisons for SARS2-RBD, SARS1-RBD, RaTG13-RBD, and NL63-RBD predicted binding sites to human ACE2 (Residues: blue highlighted occur in ^3^/_4_ RBDs, red highlighted occur in ^4^/_4_ RBDs, purple highlight is NL63 specific). **B. ACE2 Nt Distal Region**. Predicted binding sites of human ACE2 for SARS2, SARS2, RaTG13 and NL63 (blue highlighted occur in ^3^/_4_ RBDs). SARS2 residue F486 interacts with L79 and M82 whereas, N487 and Y489 serve to coordinate ACE2 protein, Nt amino region and Y83. F486, N487, Y489 are all in close proximity to disulfide Loop 4 C480-C488.

Utilizing our co-plating in vitro assay (Feinstein 2024a) and guided by the co-crystal studies of SARS2-RBD and ACE2, a set of eight mutations (Group I) targeting key residues in the Nt domain with either alanine or alteration in charge character **(Figure 2)**. Another three mutations targeted multiple residues that would coordinate ACE2 Proximal and Distal regions by N487/Y489 residues. These multiple alanine substitutions at key ACE2 residues: Q24A, Y83A; T27A, F28A, K31A, Y83A or L79A, M82A, Y83A (LMY➔A. None of the eight sets of point mutations had any effect on blocking the 16 ACE2 Nt contacts. These results seem at odds with the co-crystal data, but not with the alanine substituted RBD mutants that also failed to disrupt binding and internalization (Feinstein 2024a).

**Figure 2:**
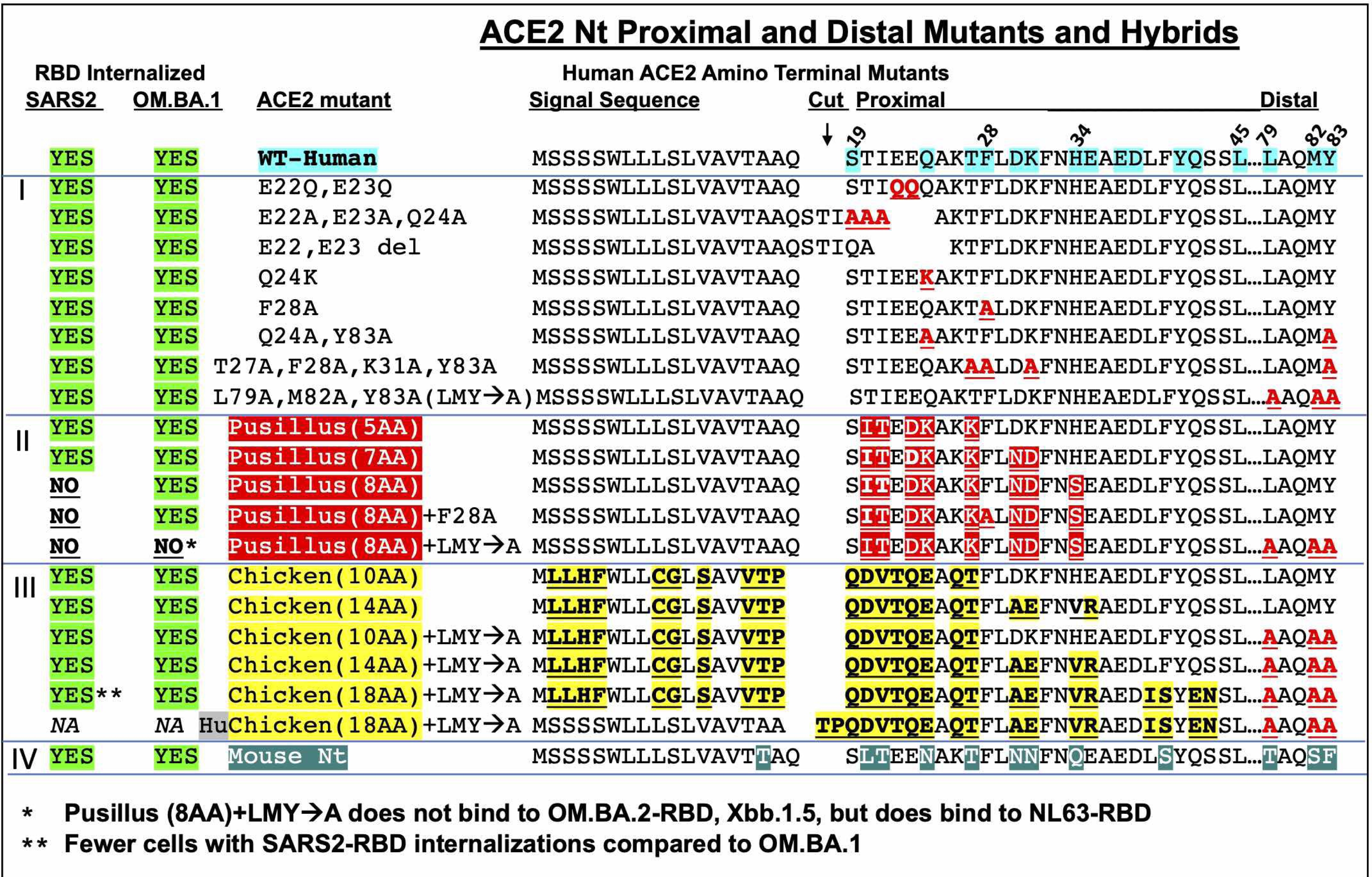
Amino Terminal human ACE2 Mutants. Twenty variants of human ACE2 (**Group I, II, III, IV**) were generated with amino terminal mutations in in the first 83 amino acids of the predicted protein. Only one mutant, HuChicken(18AA), did not show red fluorescence at the plasma membrane (PM). When RBD (green) internalization occurred, most of the red cells (ACE2 mutant) had internalized green fluorescence. To score for absence of binding (as was performed for RBD mutants), at least three touching Red/Green cells had to be observed with no green-RBD internalizations (see **Supplemental Figure S3**). **Group I mutants**: Based on crystallographic studies showing residues in the WT ACE2 (turquoise) that interact with SARS2-RBD. Mutation carrying T27A, F28A, K31A, Y83A is based on 4 residues that increase affinity of RBD to ACE2 (25) . No block of Internalization for green fluorescence for red fluorescence cells were observed in all eight variants for both SARS2 and OM.BA.1-RBD expressing cells. **Group II mutants**: Based on *R. pusillus* ACE2 not binding to SARS2-RBD (13, 14). Internalization of SARS2-RBD was observed for mutants containing the 8 amino acid substitutions, but they did not affect OM.BA.1-RBD internalization. Addition of F28A (a key residue in Y489 interaction) had no effect on OM.BA.1-RBD internalization. Only the addition of L79A, M82A and Y83A was able to abolish OM.BA.1-RBD internalization. **Group III mutants**: Based on Chicken ACE2 not binding to SARS2-RBD (13–15). In contrast *R. pusillus* mutants, none of these six mutants were able to abolish Internalization of green fluorescence in red fluorescence cells. It should be noted that NL63-RBD was internalized by both *R. pusillus 8AA* and *R. pusillus 8AA+LMY➔A* (**see Supplemental Figure 3, S3**) **Group IV mutant**: Nt Residues 1-94 are replaced with Mouse Nt residues, which have no effect on binding/internalization of SARS2-RBD or OM.BA.1-RBD.

It has been shown that ACE2 form three species R. Pusillus, Chicken, and Turkey either poorly or do not bind SARS1 or SARS2-RBD domains (13–15). This lack of binding is believed to be a distributed consequence of residues along the entirety of the ACE2 protein. But sequence analysis showed that Aves/Reptilla (**Figure 1A**) were more variable than the entire set of 408 ACE2 aligned residues. This divergence set forth a hybrid ACE2 Nt approach substituting the first 43 residues: five mutants based on R. Pusillus (Group II) and six mutants based on Chicken (Group III).

R. Pusillus ACE2 has only nine differences with the human ACE2 protein between Q19 and L45, in the Nt proximal region where SARS2-RBD is predicted to bind. Three hydrids were generated with 5 amino acids (5AA), 7AA and 8AA substitutions and tested with SARS2-RBD. Neither the 5AA nor 7AA disrupted binding/internalization. But the Pusillus 8AA completely disrupted/internalization **(Figures 2, S1)**, a furtherance of the idea that Y489A and the Loop 4 region is important for SARS2-RBD binding to Nt ACE2. All variants and mutants of SARS2-RBD failed to bind or be internalized by Pusillus 8AA: Delta, HiFrequency2020 (3Ala), HiFrequency2021(7Aala), P.B.S. 7Ala, P.B.S. 6 Ala as well as SARS1, RaTG13. **(Figure S1)**. Surprisingly, RBD-OM.BA.1 and OM.BA.2 were unaffected (**Figures 2, S1)**, as well as NL63-RBD binding/internalization consistent with its paucity of contacts with the ACE2 Nt proximal region (**Figure 1)**.

Due to the strong conservation of F28 in ACE2, in addition to its predicted interaction with Y489, an F28A mutation was added to ACE2 Pusillus 8AA hybrid to determine if OM.BA.1 or BA.2 would be blocked from binding/internalization, but no decrease in binding was observed (**Figure 1)**. These observations conclude that there is a differential reliance on the ACE Nt proximal region for SARS2 versus OM.BA.1, OM.BA.2 and NL63-RBDs, suggesting multiple modes of ACE2 binding/internalization. Subsequent RBD variants OM.BA.4 and BA.5 (same sequence) do not bind to the ACE2 Pusillus 8AA Nt mutant, suggesting that its conformation has reverted back to be more like SARS2-RBD. Finally, data from ACE2 deep mutational screening (DMS) show that SARS2-RBD-H34S does not block interactions, yet the S34H revertant in ACE2 Pusillus 7AA hybrid did rescue SARS2-RBD binding. Taken together Pusillus hybrids likely change confirmation of the Nt region, rather than disrupt any specific residue interaction between SARS-RBD and ACE2.

The proposed model for disulfide Loop 4 interactions suggested that it coordinates its binding between the ACE2 Nt proximal and distal Regions (ACE2 residues 79-83). Based on the ACE2 Pusillus 8AA mutation, SARS2 may primarily utilize the ACE2 Nt proximal region and conversely OM.BA.1 and OM.BA.2 RBDs might primarily utilize the Nt distal region. None of the Group I mutations analyzed including the Nt distal region mutant LMY➔A altered OM.BA.1 binding/internalization **(Figure 2, S2A-C)**. Data from ACE2 DMS show that SARS2-RBD-H34S has does not block interactions, but the S34H revertant in ACE2 Pusillus 7AA hybrid rescued SARS2-RBD binding. Taken together Pusillus Hybrids likely change confirmation of the Nt region, rather than disrupt any specific residue interaction between SARS-RBD and ACE2.

The RBD domains of OM.BA.1 and OM.BA.2 contain 16 mutations that differ from SARS2, which perhaps caused them to rely on both the Nt proximal and distal regions. To test this idea, the ACE2 Pusillus 8AA hybrid was combined with the LMY➔A mutant. Pusillus 8AA + LMY➔A was finally able to block OM.BA.1 and OM.BA.2 RBDs from being bond and internalized (**Figure 2, S2D-F**), confirming the hypothesis; the NL63-RBD, which does not bind to the ACE2 Nt distal regions, was unaffected as expected (**Figure S2G-I**). These Group II ACE2 mutants show that SARS2, OM.BA.1 and NL63 RBDs have differential contact points to promote their binding/internalization In contrast to the Group II hybrids, chicken ACE2 mutants (Group III) containing both the Nt proximal and distal region substitutions did not block SARS2, OMBA.2 nor NL63 RBDs from binding and internalization. Group III data suggest that some combination ACE2 Pusillus 8AA residues can block SARS2-RBD binding by a robust protein confirmational change (**Figure 2**). The totality of data for OM.BA.1 and OM.BA.2 RBD binding point to a confirmational shift in their RBDs that alters their usage for ACE2 Nt Proximal and Distal regions. Consistent with this conclusion, both Y489A and P491A are properly trafficked to the plasma membrane for the SARS2-RBD, whereas mutation in OM.BA1-RBD fail to traffic to the plasma membrane (Feinstein 2024a).

Mice cannot be infected by SARS2, which is why mutant mice have been made generated with the human ACE2 CDS replacing the mouse ACE2 gene. A third hybrid (Group IV) with mACE Nt proximal and distal was generated with 25 of the first 94 residues altered (**Figures 2, S3**).

This Group IV hybrid contained mutations at similar residues to the Pusillus 8AA as well as an LMY➔TSF mutation. Yet, no alteration in binding or internalization was observed for SARS2, OM.BA.1 and NL63 RBDs. Thus, substantial changes in the ACE2 Nt proximal and distal regions suggests a lack of specificity for RBD binding. We confirmed and quantified the critical results for Pusillus 8AA, Pusillus 8AA +LMY➔A, and mACE2 Nt 1-94 by our Dissociation-Plate-FACs *(DPF) assay* (Feinstein 2024a) with RBDs from SARS2, OM.BA.1, NL63 and a newer variant Xbb.1.5.

Indeed, FACs quantification of double positives for SARS2, OM.BA.1, NL63 mimicked the co-plating assay results with Xbb.1.5 showing binding and internalization like that of OM.BA.1 in all four Nt ACE2 hybrids (**Figure 3, Supplemental Data File S2**). In contrast to OM.BA.1, Xbb.1.5 did not cause a change in fluorescence to Pusillus 8AA or Pusillus 8AA +LMY➔A-Cherry cells. OM.BA.1 despite a lack of robust binding to Pusillus 8AA +LMY➔A (8.75±1.94), still manage a modest reduction to its fluorescence (17.2%±2.2%). Finally, a fourth mutant mACE 1-357 was generated, which abolished binding for RBDs SARS2(1.45±1.37), OM.BA.1(5.71±0.89), and Xbb.1.5(3.98±2.08), and but not for NL63(26.7±1.60) (**Figure 3**). An interaction could be observed for OM.BA.1 by the 24.8%±0.04% loss of ACE2 fluorescence. These data are consistent with NL63 having a greater set of contacts with residues 321-330 of ACE2 (**Figure 4**).

**Figure 3:**
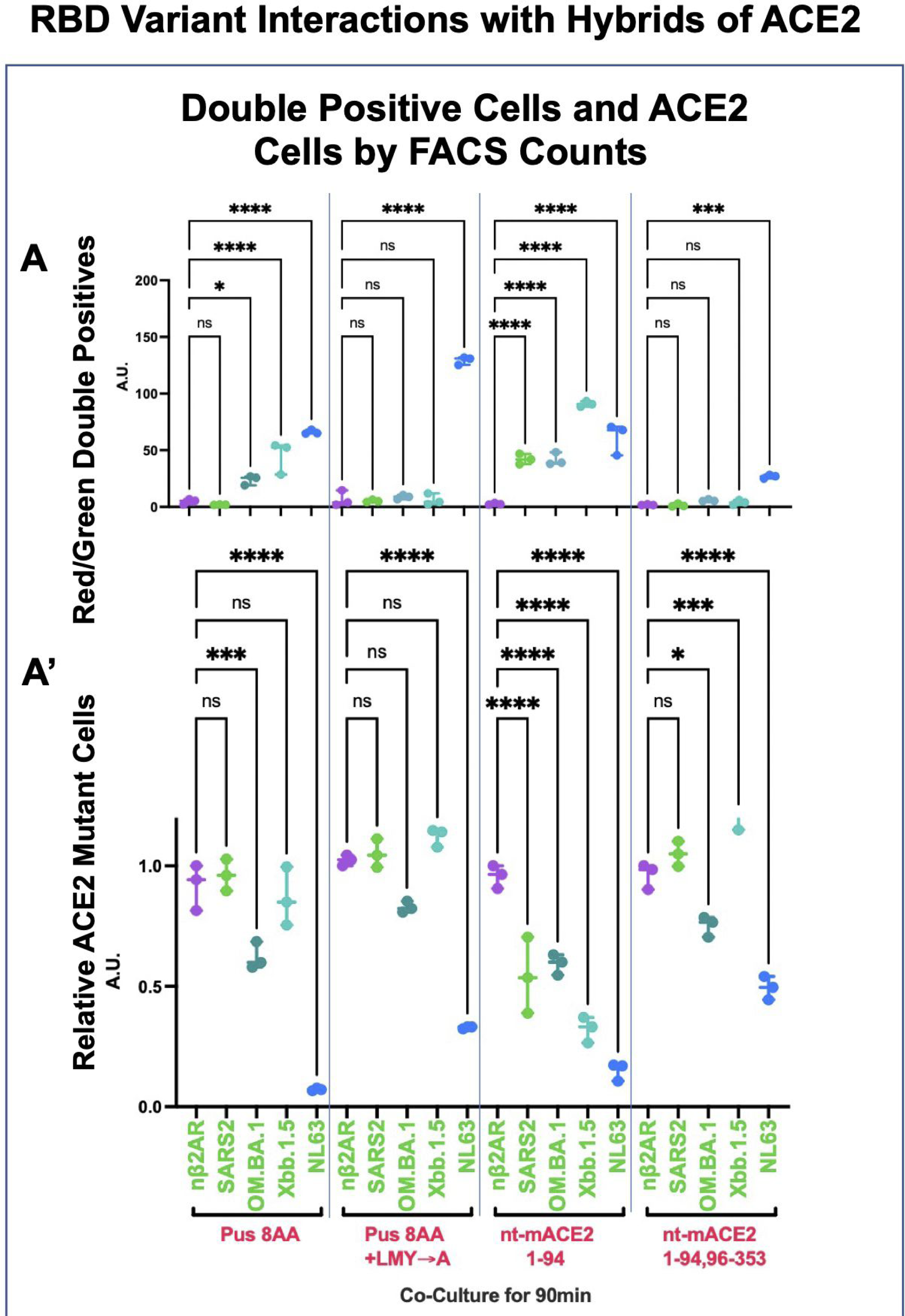
RBD Variants interactions with Mutated ACE2. **A.** Results from visual inspection for Pus8AA, Pus8AA+LMY➔A and Nt-Mouse residues 1-94 (Figure 2) were further quantified by the *DPF assay* for nβ2AR, SARS2-RBD, OM.BA.1-RBD, Xbb.1.5-RBD and NL63-RBD. In addition, a near complete swap of human residues 1-353 with equivalent region of mouse residues. NL63-RBD show double fluorescence in all four ACE2 mutants and a reduction of ACE2 fluorescence, revealing all ACE2 constructs have the capacity to function. SARS2 Variants show differences (**see Supplemental Figure S3 and Data File S2**). *For Pus8AA*: only SARS2-RBD shows no binding and hence no double fluorescence (1.92±0.17). *For Pus8AA+LMY➔A*: SARS2-RBD, OM.BA.1-RBD, show no binding (5.23±0.99) and (8.75±1.94), respectively. *For Nt-Mouse residues 1-94*: All variants show binding/double fluorescence (58.22±28.20). *For Nt-Mouse 1-94,96-353:* All variants show **no** binding/double fluorescence (3.71±2.14). **A’**. The ACE2 levels showed variability for Xbb.1.5-RBD as it does not down regulate Pus8AA(a loss of 13.3%±12.1%) as we previously showed for ACE2 (Feinstein 2024a) but does down regulate the Nt-Mouse residues 1-94 ACE2 chimera(a loss of 67.8%±5.3%). To date, Xbb.1.5 is the only variant where binding **can be** decorrelated from an ACE2 decrease in level of expression.

**Figure 4:**
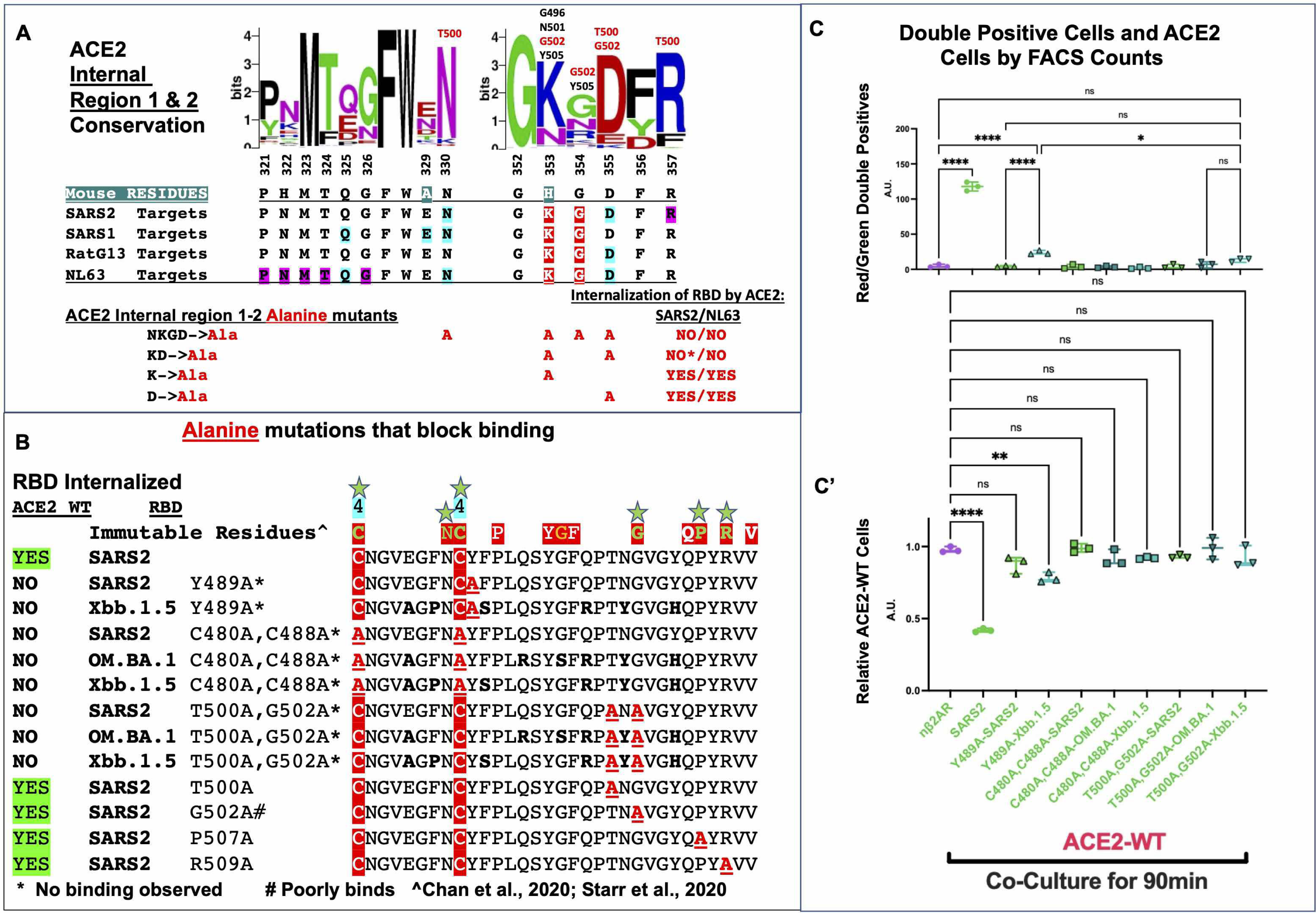
Internal human ACE2 mutants and RBD interaction mutants. **A.** ACE2 Internal Region. Predicted binding sites of human ACE2 for SARS2, SARS2, RaTG13 and NL63 (Residues: blue highlighted occur in ^3^/_4_ RBDs, red highlighted occur in ^4^/_4_ RBDs, purple highlight is SARS2 specific). ACE2 mutants tested based on conserved predicted binding sites: 1) N330A, K353A, G354A, D355A (NKGD➔A); 2) K353A, D355A (KD➔A); 3) K353A; 4) D355A. Both SARS2 and NL63 RBDs were tested in two ways: cotransfection or separate transfections followed by overnight co-plating (**see Supplemental Figure S5**); Co-transfection of RBD-β2AR-GFP and ACE2-Cherry that could interact typically internalized membrane distribution of both proteins. Mutant NKGD➔A, which poorly traffics to the plasma membrane, shows no binding to either SARS2 or NL63 RBDs, KD➔A shows no binding to NL63 RBD, but some binding to SARS2-RBD without internalization. By contrast the single mutants K353A and D355A mutants both revealed binding and internalization of both NL63 and SARS2 RBDs. **B.** Alignment of different SARS2/OM.BA.1/Xbb.1.5 mutants and that were tested for block binding (**see Supplemental Figure S4**); Region consists of residues 480-511 from SARS2. Bolded residues differ from SARS2. Bolded Red and underlined residues are alanine mutations. Mutants tested were Y489A and the double mutant C480A, C488A on SARS2, OM.BA.1, Xbb.1.5 that previously showed no binding (Feinstein 2024a). In addition, the T500A, G502A double mutant, which is predicted to bind to KGD region of human ACE2. Visual observations were noted and quantified. **C.** *DPF assay* quantification for mutants that did not predict binding by visual inspection. Only Xbb.1.5-Y489A-RBD showed slight binding as some double positive Red/Green cells could be observed (23.67±2.87) and some loss of ACE2-Cherry fluorescence is shown (21.7%±3.4%).

### Cotransfection of membrane tethered RBDs and ACE2

In vitro co-plating while simple, requires visual inspection of RBD expressing cell touching an ACE2 expressing cell. This assay is robust but can be difficult to quantify weak binding without internalization. The *DPF assay* is very quantitative, but more labor intensive. To more quickly rule out weak interactions, a co-transfection and plating assay was set up to see if protein interactions could be observed within the same cells as is reminiscent of the interactions published decades ago by the SEVENLESS-BOSS (16) proteins as well as the NOTCH-DELTA (17) proteins. Co-transfection between nβ2AR-GFP with ACE2-Cherry in the same cells and imaged after 16hrs reveals robust membrane trafficking for both proteins in the same cells (**Figures S4A-A’’**). By contrast, co-transfection of SARS2-RBD with ACE2-Cherry show robust SARS2-RBD at the membrane and filopodia, and all ACE2-Cherry internalized (**Figures SB-B’’)**. Internalization of ACE2 by RBD by intra- or intercellular interactions may be through clathrin (18);. This final piece of evidence firmly concludes that ACE2-Cherry is subordinate to the effects of the SARS2-RBD protein and does not cause cell death. This assay confirms and extend results observed on ACE2-Cherry membrane localization in Feinstein 2024a: Y489A (**Figure S4C-C’’**) and P.B.S 7Ala (**Figure S4D-D’’**) have no effect, C480A, C488A (**Figures S4E-E’’**) had little effects and P.B.S. 6Ala (**Figures G-G’’**) completely internalized it. Consistent with co-plating experiments, co-transfection of SARS2 and OM.BA.5 RBDs with Pusillus 8AA hybrid ACE2 did not show binding/internalization (**Figures S5A-A’’, B-B’’**) whereas co-transfection with OM.BA.1 RBD did show internalization (**Figures S5C-C’’**). Surprisingly, OM.BA.1 RBD did internalize Pusillus 8AA +LMY➔A (**Figures S5D-D’’**), despite it not forming double fluorescent cells by *DPF assay* (**Figure 3**), but perhaps consistent with the loss of fluorescence (**Figure 3**). Co-transfection proved an effective way for simultaneously assessing membrane trafficking of ACE2 and RBD mutants as well as determining if proteins could interact.

### ACE2 Internal region mutagenesis

Interpretation of the data thus far suggests the ACE2 Nt proximal region is necessary for SARS2-RBD binding/internalization and both ACE2 Nt proximal and distal regions are necessary for OM.BA.1 and BA.2-RBDs; Neither region is necessary for NL63-RBD binding/internalization. SARS2-, SARS1-, RaTG13- and NL63-ACE2 co-crystals map residues to two additional Internal regions of ACE2: residues 321-326 (PNMTQG); 330 (N) and 353-356 (KGDF), with NL63 contacting all of these residues; with SARS2, SARS1 and RaTG13 contacting different subsets (**Figure 4A**). All four of these RBDs contact ACE2 conserved residues K353 and G354 (**Figure 4A**). G354 contact is not likely as important as the D355 contacts found in three RBDs (*not* SARS1). NL63-RBD being unaffected by the Pusillus 8AA+LMY➔A mutant nor mACE2 1-357 suggests that this Internal region is key to triggering binding and internalization of all RBDs. It was previously shown that ACE2 mutants K353A and D353A nearly abolish all SARS and NL63 infections (19–21). Several alanine substitutions tested this region for interfering with SARS2 or NL63 binding/internalization: quadruple mutant-N330A, K353A, G354A, D355A, the K353A and D355A pair, and individual mutations in K353A and D355A (**Figure 4A**). The NKGD➔A and KD➔A mutants blocked both SARS2 and NL63-RBD binding and internalization (**Figures 4A, S5E-E’’, S5F-F’’**). But the individual K353A and D355A mutations had no effect (**Figures 4A**). The internal region mutants point to a common binding site on ACE2 that all RBDs utilize for binding/internalization.

Co-crystal studies of RBD and ACE2 suggest K353-G354-D355 in ACE2 interacts predominantly with T500 and G502 in the SARS2-RBD, a region distinct from the Loop 4 region. G502, P507, and R509 all represented immutable residues from DMS studies suggesting they could all be key residues for ACE2 interaction. Co-transfections were carried out for this region of SARS2-RBD: T500A, G502A, P507A, P509A and T500A, G502A double mutant (**Figures 4B, S4H-L’’**). Only the double mutant T500A, G502A was able to completely abolish ACE2 binding (**Figures S4J-J’’**). These data argue that two regions of SARS2-RBD work tother to coordinate binding to ACE2: T500 and G502 coordinate the internal region of ACE2 while Y489 and the Loop 4 disulfide bond (C480-C488) coordinate the Nt distal and proximal residues. Perhaps the effect of these mutations is not universal across variants as the Y489A mutation in OM.BA.1 disrupted membrane trafficking, likely due to a disruption in RBD folding. The *DPF assay* was once again diploid to compare these three types of mutations in the SARS2, Xbb.1.5 and OM.BA.1 RBDs (**Figures 4C, C’**); Both C480A, C488A(3.04±1.04) and T500A,G502A (7.87±4.80) disrupted binding/internalization (double positives) of ACE2 across all variants (**Figures 4C, C’**). The sum total of all RBD mutations we produced suggest binding is localized to a stretch of residues between Y489-G502, which is also where blocking antibodies target (22).

### Can RBD binding to ACE2 be antagonized by KGD containing peptides?

A common binding site on ACE2 for both NL63 and SARS2-RBDs suggested that a competitive antagonist based on the Internal region 2 amino acid sequence could be derived. Six peptides were generated and tested for functional antagonism in the solution binding assay at μM concentrations (**Figure S6A**). None of the peptides interfered with RBD-ACE2 binding for both NL63 and SARS2-RBDs. However, one of the peptides (GKGD) caused red fluorescence of ACE2-Cherry cells to surround SARS2-RBD expressing cells (**Figure S6A insets**). Because the tripeptide KGD is similar to the high affinity RGD tripeptide for fibronectin (23), two RGD containing peptides for blocking activity were also tested at μM concentrations, but neither had any impact. Next, the GKGD peptide alone was tested in the *DPF assay* with a preincubation of 100μM GKGD peptide, followed by FACs analysis for both SARS2 and NL63-RBDS (**Figure S6B,B’**); Again, no mitigation of SARS2-ACE binding was observed. Finally, purified His-tagged SARS2-RBD (60ng) (Feinstein 2024a) was added to ACE2 cells with or without preincubation with 49μg of the GKDG peptide in the *DPF assay*; No decrease in RBD binding was observed (**Figures S6C-C’’**). Taken together, these data argue that small KGD containing peptides do not bind RBDs at high enough affinity to block ACE2 binding.

## Discussion

### The disulfide Loop 4 region interacts with Nt proximal and distal regions of ACE2

The importance of disulfide Loop 4 located residues F486, N487 and Y489 from the Feinstein 2024a analysis, would seem to correlate with the Nt region in ACE2 as critical for binding. RBD residues 486-489 bridge Nt proximal region residues Q24, T27, F28 with Nt distal region residues L79, M82, Y83 (LMY) (**See Figure 1B**). But ACE2 mutations (Group I) targeting these residues had no effect on RBD binding and internalization. Group II Nt Hybrids with R. *Pusillus* sequences abolished SARS2-RBD interactions, but not OM.BA.1. Combining Nt proximal R.Pusillus residues and Nt distal substitutions LMY➔A did lead to a complete block in binding and internalization by OM.BA.1-RBD. These results suggest that the 15 amino acid differences in OM.BA.1-RBD lead to a new protein conformation, which is consistent with the absence of Y489A protein trafficking in a OM.BA.1-RBD. At first approximation, these data argue that Nt proximal and distal ACE2 regions are necessary for binding and internalization of SARS2 and OM.BA.1-RBDs. But the Nt mACE 1-94 hybrid still providing binding and internalization suggest a lack of specificity of action.

### SARS-CoV-2 and H-CoV-NL63 both require common Internal ACE2 region for binding

The NL63-RBD binding and internalization by ACE2 was unaffected by any of the 20 Group I/II/III/IV Nt proximal or distal mutations. Those results are consistent with the co-crystals showing only a few contacts by NL63-RBD in the Nt proximal and distal domains. By contrast, NL63-RBD has extensive contacts in both Internal region 1 and 2 where as SARS2-RBD has the majority of its contacts to Internal region 2. Previously it was shown that K353A had substantial impact on both NL63 and SARS2-RBD binding in pseudovirus infections but did not abolish it. Our co-plating and co-transfection assays also did not reveal complete blocks in binding and internalization. While the single substitution D355A did not have an impact, the double mutant KDèA abolished NL63 binding and blocked internalization of SARS2-RBD. The internal region represents a second ACE2 site, when mutated, abolishes nM SARS2-RBD binding, consistent with the prediction that high affinity binding is distributed.

Based on the RBD mutants identified here and in Feinstein 2024a, it appears the most critical RBD region for interaction with ACE2 may be further refined to a 23-residue region from C480 to G502, which encompasses the Loop 4 disulfide bond (C480-C488). It is noteworthy that this region represents the highest density of residue contacts specific to the m396 neutralizing antibody against SARS2 (22, 24). The disruption of RBD variant binding with the double mutants C480A, C488A and T500A, G502A suggest this region remains critical for any new variant binding to ACE2. It is possible that a peptide inhibitor utilizing this 23-residue stretch could be the basis of a competitive or allosteric inhibitors.

A model emerges (**See Figure 5**) whereby the Internal region 2 may be critically important for stable RBD-ACE2 binding/internalizations but require secondary contacts for stabilization. In the case of NL63-RBD, those additional contacts are in the Internal region 1 and for SARS-RBDs, those additional contacts are in the Nt proximal and distal regions. It still may be possible to identify a high affinity agonist to RBD based on the NL63/SARS2 target KGD motif in ACE2 that could mitigate binding. Finally, peptides against both the Nt and Internal region binding sites might work synergistically to inhibit the spread of SARS-CoV-2.

**Figure 5:**
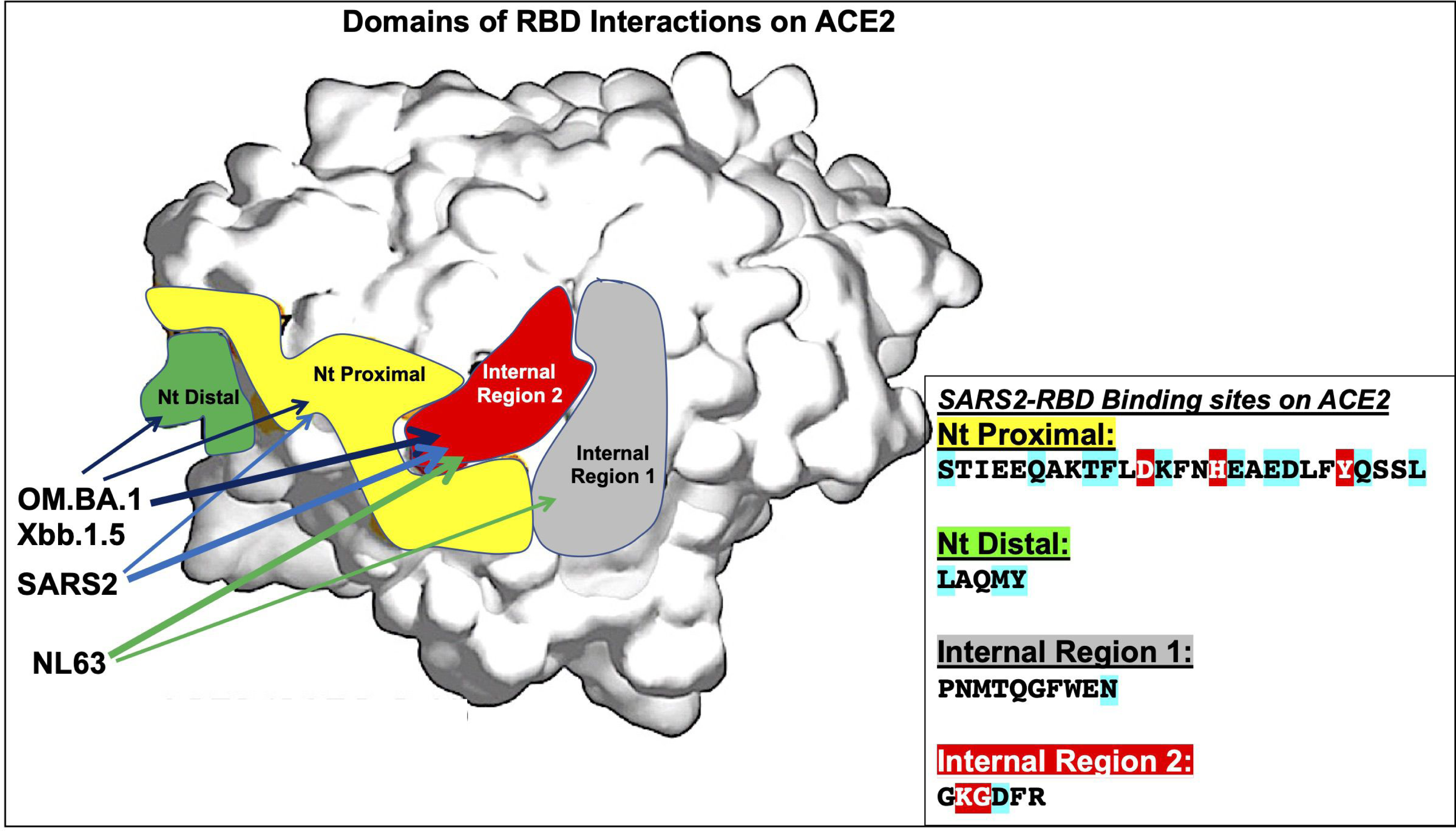
Proposed interpretation of RBD interactions on ACE2 structure. Space filling model of ACE2 structure (26) modified by highlighting Nt proximal (yellow), Nt distal (green), and Internal regions 1 (gray) and 2 (red). SARS2-RBD does not bind/internalize if *either* Nt proximal or Internal region 2 (K353, D355) are mutated. OM.BA.1 and OM.BA2-RBDs are not bond/internalized if *either* mutation in both Nt proximal and distal ACE2 or Internal region 2 (K353, D355). NL63-RBD requires Internal region 2 (K353 and D355) for both binding/internalization. Inset shows **SARS2-RBD** interactions within the four domains (**see** Figures 1**, 4**) (residues with blue highlights are predicted binding sites that occur in ¾ of SARS2, SARS1, RaTG13, and NL63-RBDs; residues with red highlights are predicted binding sites in all 4 RBDs). The RBD interaction data taken together suggests that Internal region 2 binding for SARS1/SARS2/RaTG13 is weak and requires Nt proximal and distal contacts for stabilization (dark blue, light blue arrows), whereas NL63 has additional contacts in the Internal region 1 (green arrows) that supports binding to Internal region 2 (thick arrows). The ability of Y489A to nearly abolish SARS2-RBD binding to ACE2 suggests that none of the domains by themselves have strong binding to ACE2. Overall, the data suggests that Internal region 2 is critical for internalizations of RBDs.

### Implications

*Internal regions of ACE2:* The published NL63-RBD affinities to ACE2 are similar those of SARS2, but likely utilize internal regions of the protein and points to this as the key region for binding/internalization of ACE2. It is worth noting that NL63-RBD is more robust at reducing ACE2 fluorescence (Feinstein 2024a and **Figure 3A’**), which also suggests that the better a protein binds to the internal ACE2 region, then the more robust internalization. The changing nature of SARS2-RBD binding to ACE2 argues that variants could emerge that rely more on the ACE2 internal region for binding like NL63-RBD. It is unclear how COVID-19 disease progression would occur if ACE2 were to be more rapidly internalized.

An antibody or peptide blocking strategy targeting the KGD region on ACE2 in the nasal epithelium could radically reduce the ability of SARS2-CoV-2 to infect sustentacular cells. Mitigating viral amplification in the nasal cavity will likely lead to reduced viral particles, fewer symptoms, and most importantly less asymptomatic spread, which could lead to an improved prognosis for the current and future pandemics.

## Materials and Methods

See Feinstein 2024a for constructs (**Data File S1)** and materials and methods.

Calculations and graphs were made in GraphPad Prism. Mean and standard deviations were derived in Excel; See Feinstein 2024a (**Data File S2)** for details.

Peptides synthesized from GenScript. RGD purchased from R&D systems, #7723/10mg. Cilengitide purchased from Tocris, #5870.

## Acknowledgements

Special thanks to Masayo Omura for providing logistical and intellectual support that helped drive the success of these experiments. Many thanks to Masayo Omura, Ivan Rodriguez, Sergio Bernal-Garcia and Thomas Bozza for advice on the experimental design and interpretation of results; Mary Slavinsky and Masayo Omura for technical support in plasmid production; and Sergio Bernal -Garcia for help with GraphPad Prism and Fiji. Thanks to Ivan Rodriguez, Vince Racaniello, Masayo Omura, and Sergio Bernal Garcia for providing feedback on the manuscript. Tomas Baumgartner Manager, Flow Cytometry Core Facility, Weill Cornell Medicine for support on flow cytometry of single cells. Many thanks to Delia Tomoiaga, Claude Rose Levine and Mark Levine for supporting my efforts to design novel experiments that might help tackle the pandemic of our lifetime. This work was supported by Hunter College and NIH grants# NIH R01 NS091439-01 and NIH R01 DC020764-01.

## Supplemental information inventory

**Figure S1:**
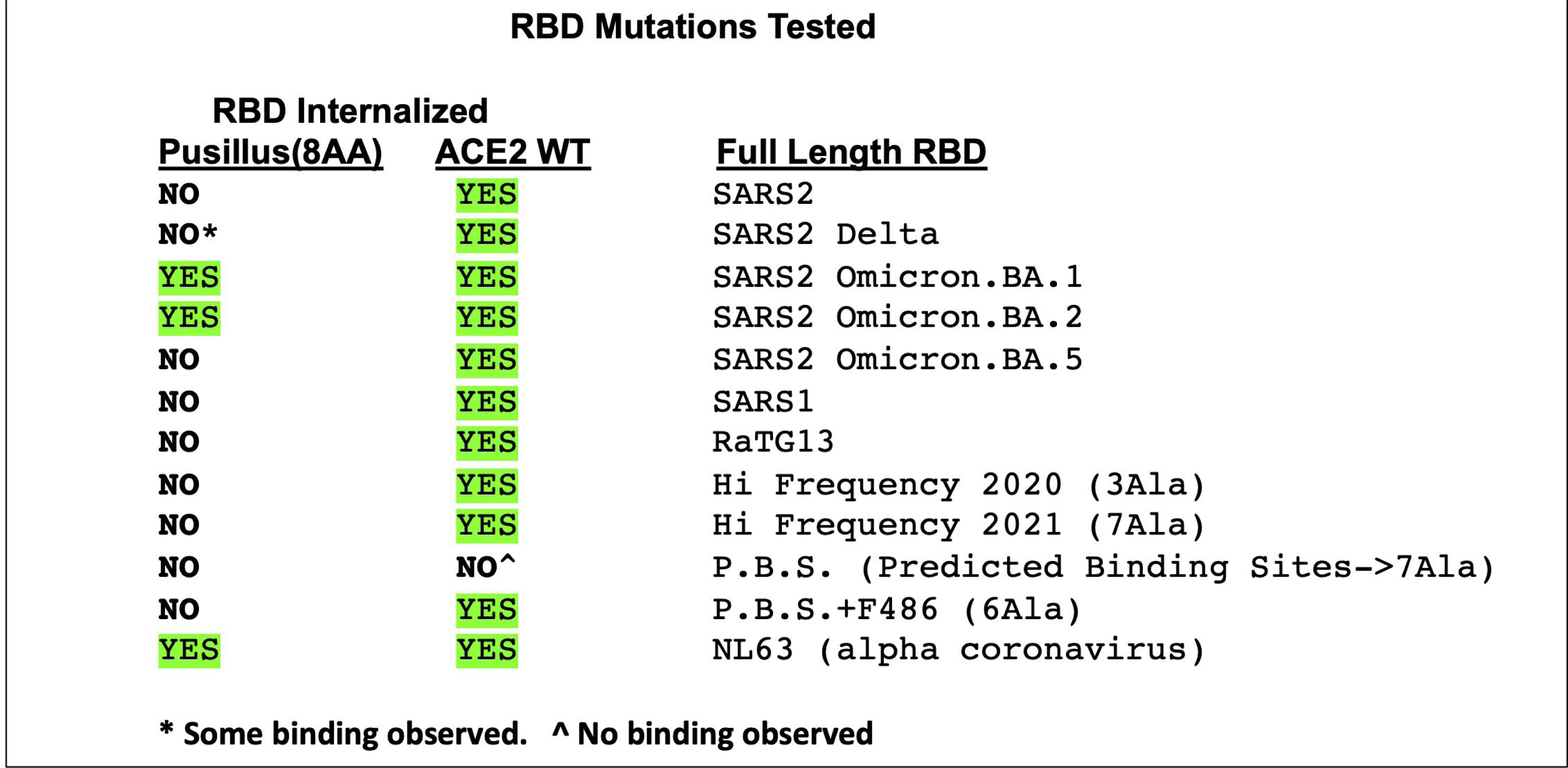
RBD mutations tested with Pus8AA+LMY➔A-ACE2. Co-plating RBD constructs with Pus8A-ACE2-Cherry cells. A broader set of RBD mutants and variants of SARS2 are tested by visual inspection **(see also** Figure 2). Only OM.BA.1-RBD, OM.BA.2-RBD, and NL63-RBD can robustly bind to Pus8AA-ACE2-Cherry. All RBD mutants that could bind to ACE2-Cherry, did not bind to the Pus8AA-ACE2-Cherry.

**Figure S2:**
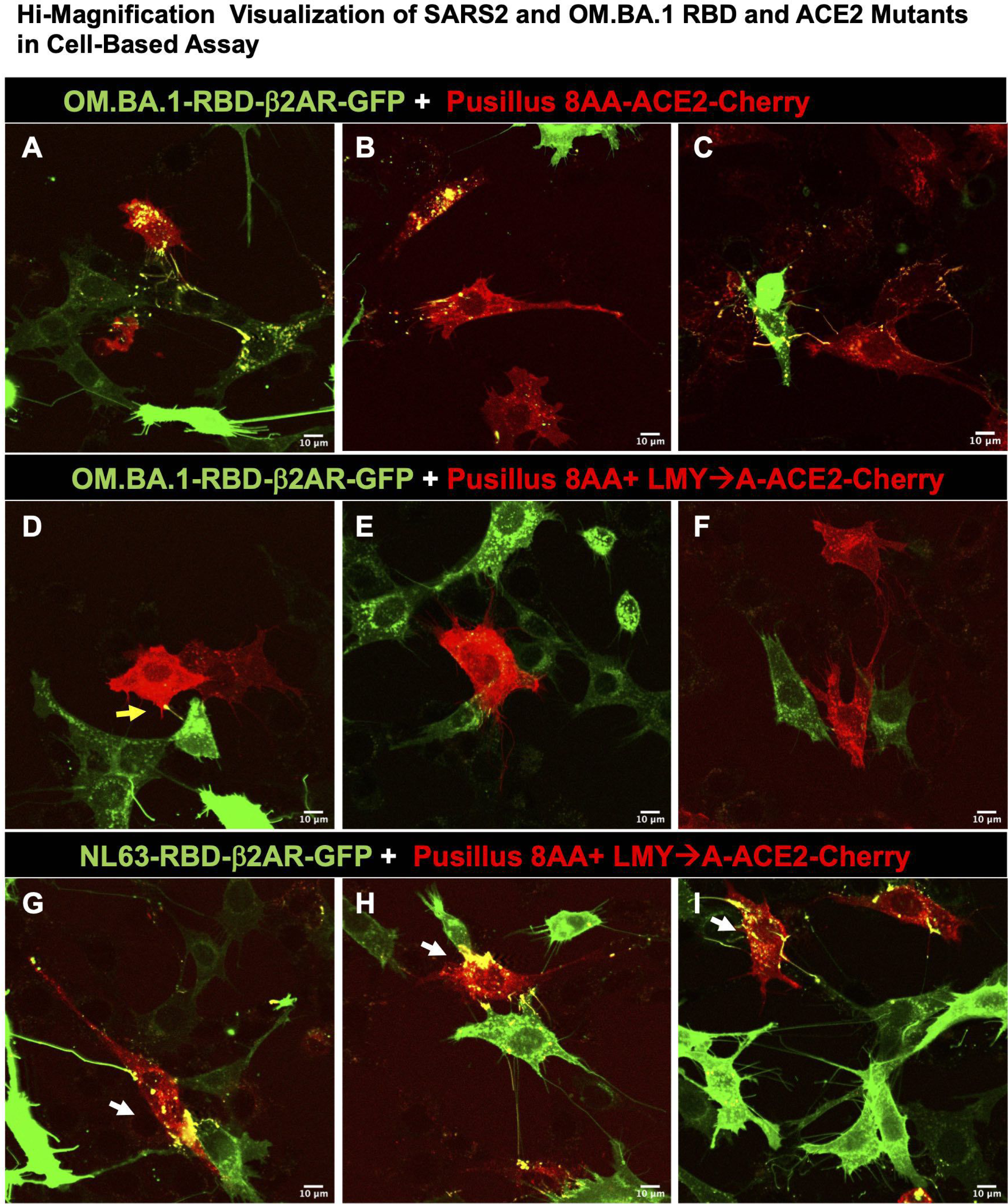
Visualization of OMBA.1 RBD mutants and NL63-RBD in Cell-Based assay with ACE2 mutants. **A-I.** Confocal images for co-plated RBD variants and mutants with ACE2-Cherry cells. **A-C**. Three representative images showing OM.BA.1-RBD protein readily internalized into Pus8AA-ACE2 express cells. **D-F.** Three representative images showing OM.BA.1-RBD protein not internalized by Pus8AA+LMY➔A-ACE2 expressing cells. **G-I.** In contrast, three representative images showing NL63-RBD protein readily internalized by Pus8AA+LMY➔A-ACE2 expressing cells.

**Figure S3:**
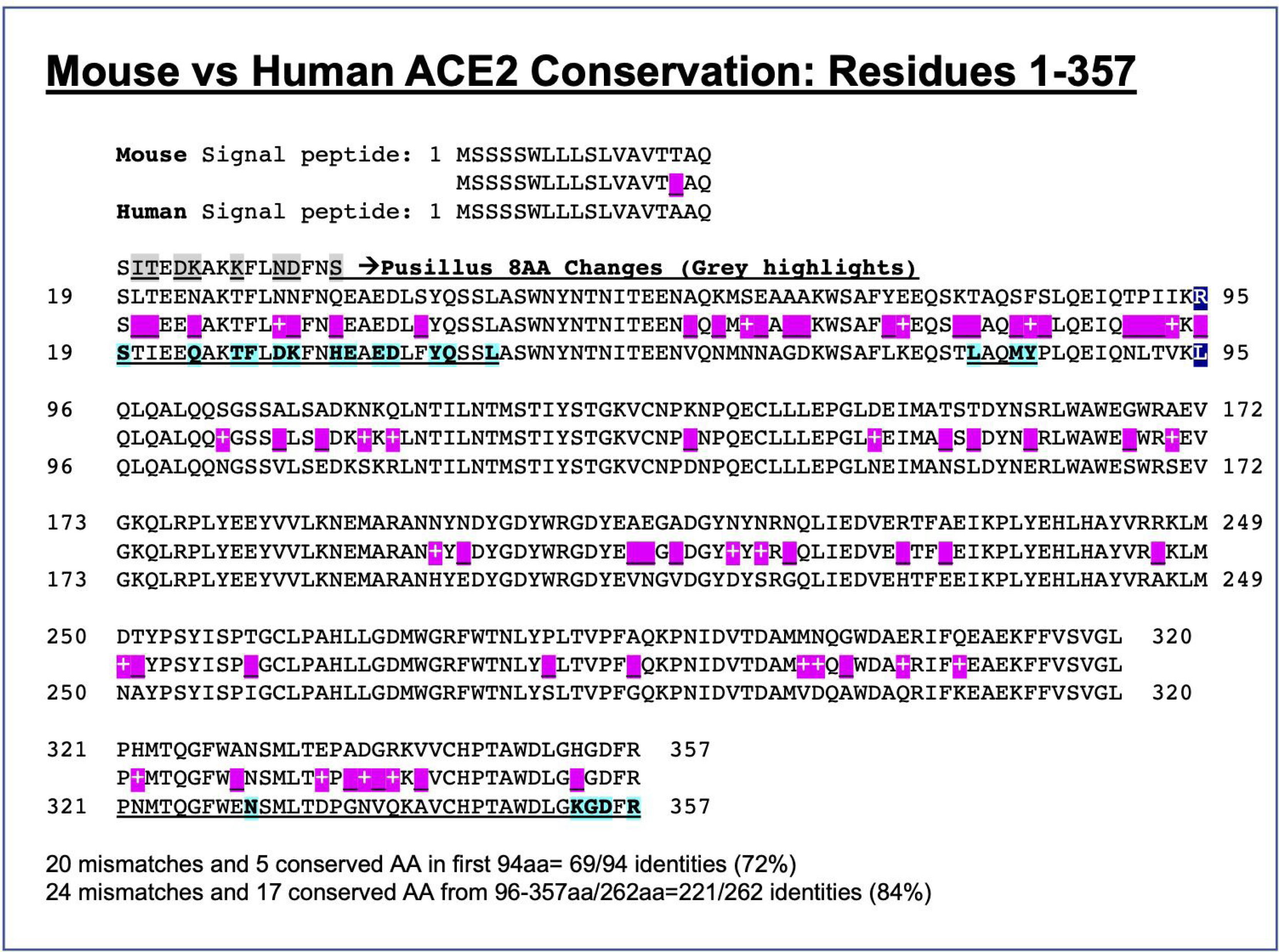
Mouse vs Human ACE2 among first 357 residues. First 357 ACE2 residues compared between mouse and human shows 87% identity. Consensus sequence shows 45 mismatches of which 22 are conserved (purple shading). Human residues predicted to interact with SARS2-RBD are shaded turquoise and bold. Nt-Proximal region (residues 19-45), Nt Distal (residues 79-83), and Internal regions 1 & 2 (residues 321-357) are underlined.

**Figure S4:**
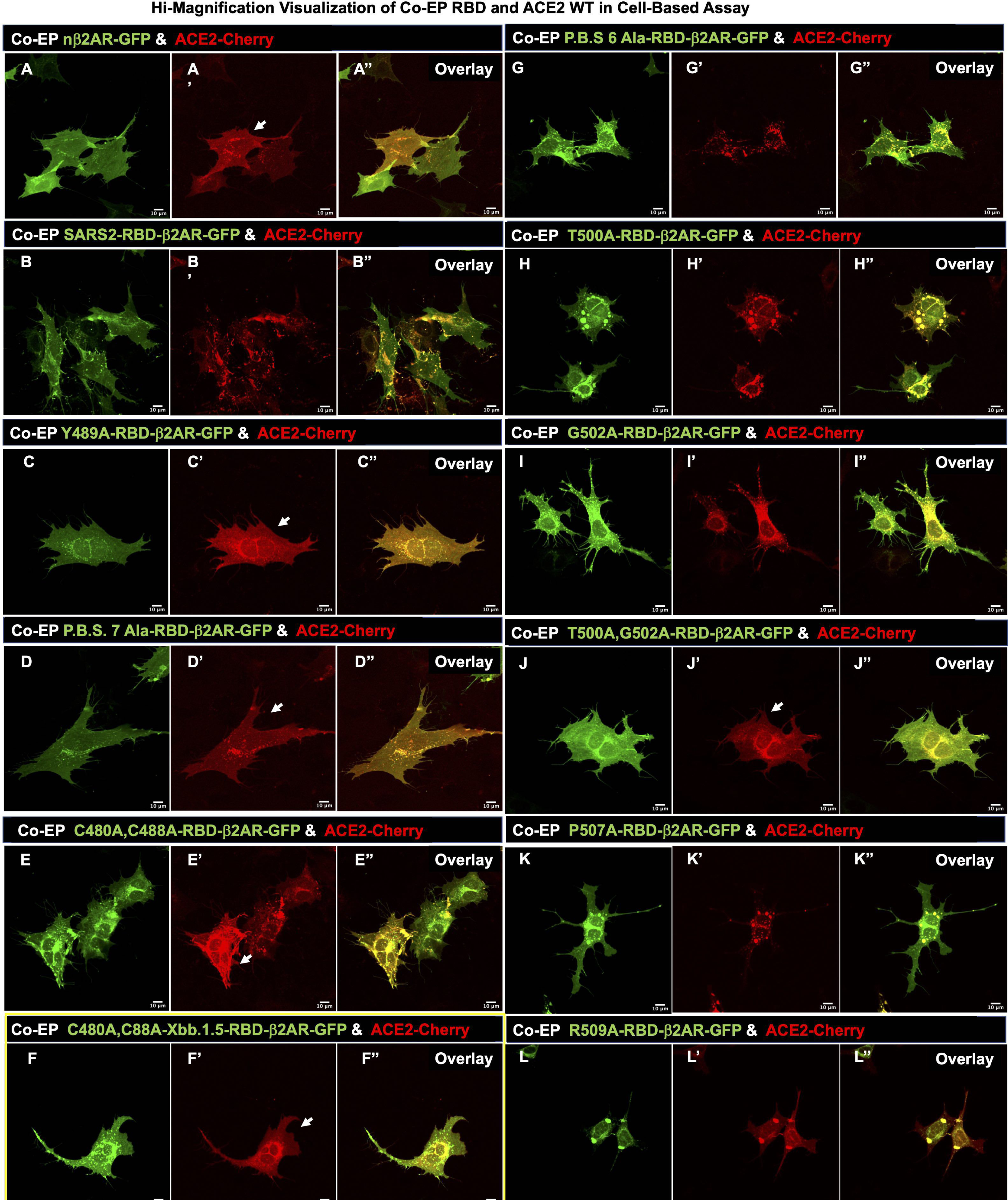
Co-EP with RBD mutations and ACE2 A-L, A’-L’, A’’-L’’. Confocal images for Co-EP of GFP-tagged RBD variants and mutants and controls along with ACE2-Cherry, imaged after 16hrs. Green **(A-L)** and Red **(A’-L’)** channels have been split, and overlay (**A’’-L’’**). **A-A’’**. β2AR and ACE2 show uniform membrane expression. **B-B’’.** SARS2-RBD and ACE2 show internalization of all ACE2-Cherry fluorescence. **C-L, C’-L’, C’’-L’’.** Of the four assays (DPF assay; Cell-Dissociation assay; Co-plating assay; Co-EP RBD/ACE2 assay) that were set up, the Co-EP proved to be the most sensitive way to determine any latent binding between RBD and ACE2 proteins. **A’, C’, D’, F’, and J’** show no alteration of membrane ACE2 expression. **B’, G’, H’, K’, L’** show robust internalization of ACE2-Cherry. **E’, I’** show only modest internalization of ACE2-Cherry.

**Figure S5:**
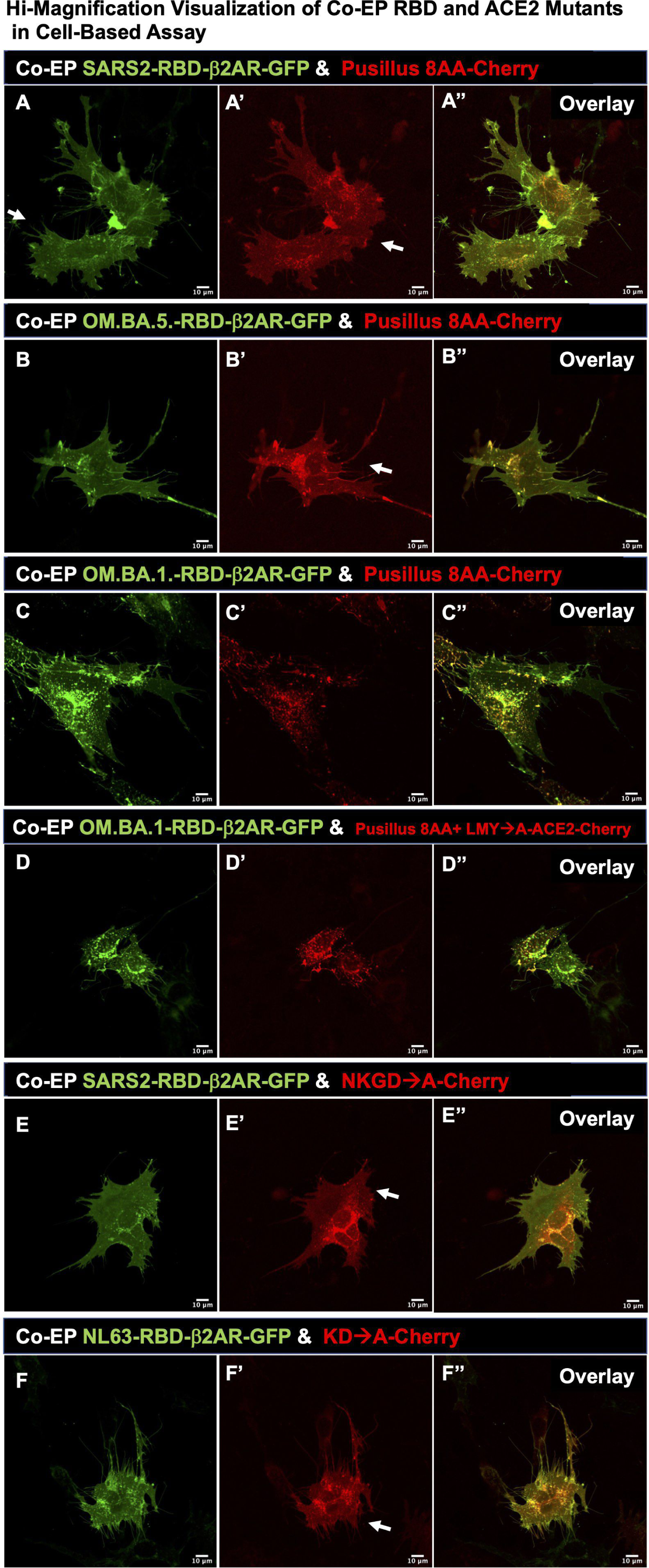
Co-EP with RBD variants and ACE2 mutants A-F, A’-F’, A’’-F’’. Confocal images for Co-EP of GFP-tagged RBD variants and mutants and controls along with ACE2-Cherry, imaged after 16hrs. Green **(A-F)** and Red **(A’-F’)** channels have been split, and overlay (**A’’-F’’**). Omicron.BA.1 clearly shows differential binding to Pusilus 8AA-Cherry compared to SARS2 and OM.BA.5-RBDs. OM.BA.1-RBD shows some internalization of ACE2-Cherry (**C’**), which shows the sensitivity of binding in this assay compared to co-plating assay in **Figure S3 A-F** and DPF assay in Figure 3. Both SARS2 and NL63-RBDs show dependence on Internal Region 2 for binding ACE2-Cherry with no alteration of membrane ACE2 expression (**E’, F’**).

**Figure S6:**
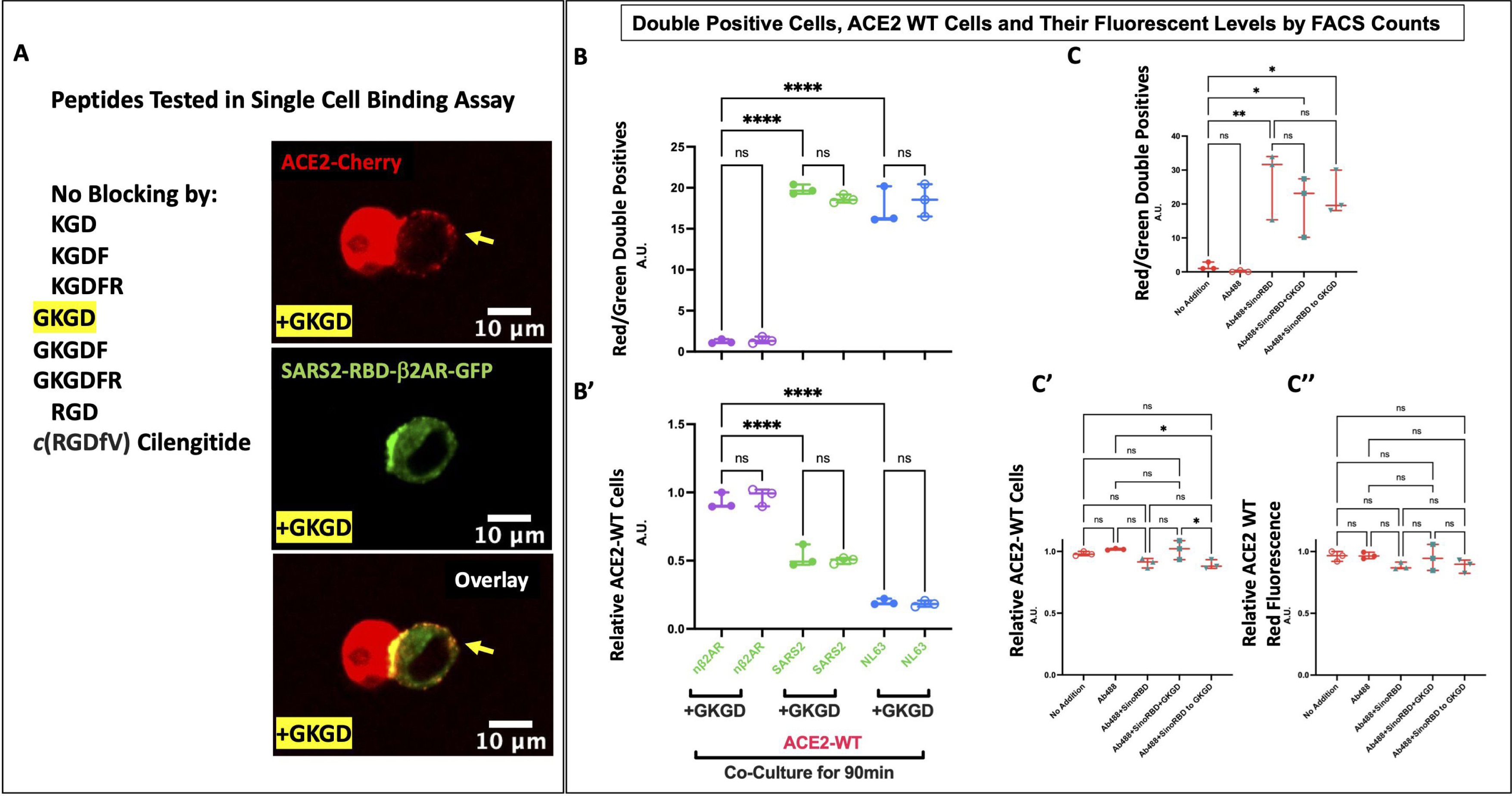
KGD containing peptides do not antagonize RBD binding to ACE2-Cherry. **A**. The necessity of K353-G354-D355 sequence in ACE2 suggested that KGD derivative peptides could block RBD binding. Six peptide versions of KGD and two RGD containing peptides were tested for their ability to block SARS2 or NL63 RBDs from binding ACE2-Cherry in the 30min dissociated cell-based assay. Greater than 100 micromolar concentrations for all 8 peptides did not block binding. However, one peptide GKGD (yellow highlight and inset) appeared to cause a phenotype that was not previously observe: ACE2 protein being redistributed on the adjoining SARS2-RBD cells. **B and B’**. *DPF assay*: analysis of Co-plating GFP and Cherry expressing cells after 90min. Data shows that double florescent cells are found in both SARS2 and NL63 irrespective of preincubation with 120μM GKGD peptide with subsequent 90min incubation at a 1:20 dilution (6μM GKDG). Concomitant loss of ACE2-Cherry expression was observed in both ACE2 cells in all RBD conditions whether or not the peptide was present. **C-C’’**. *DPF assay*: **C**. Double fluorescence after 90minute incubation of ACE2-Cherry cells with Ab488 (1μg/4mls of Goat-anti-HIS-Tag-Alexa488 Goat antibody [R&D#: IC050G]) with and without SinoRBD (60ng/4mls of 2019-nCoV-HisTag-RBD [Cat#40592-V08H; 234 residues, R319-F541]). 1000x-GKGD peptide (49μg) 10min pre-incubated with 1μg Ab488 + 60ng SinoRBD in 200μls PBS, then added to 4mls of media or GKDG added to media (49μg/4mls) plus 200μls PBS with 10min preincubated of 1ug Ab488 + 60ng SinoRBD. No reduction of double fluorescence is observed when GKGD peptide is added. **C’** ACE2-Cherry cells with Ab488, Ab488+SinoRBD, or GKGD+Ab488+SinoRBD show no changes in fluorescence. **C’’**. High expressing ACE2-Cherry fluorescence cells also show no changes in fluorescence in all conditions.

